# High-efficiency, site-specific integration of kilobase-scale DNA into plant genomic safe harbors via PrimeStack editors

**DOI:** 10.64898/2026.04.13.718181

**Authors:** Edith Sánchez, Khalid Sedeek, Haroon Butt, Magdy Mahfouz

**Author notes:** **Corresponding author**: Magdy M. Mahfouz.

## Abstract

Precise, site-specific integration of large DNA sequences into plant genomes is a cornerstone of crop biotechnology and synthetic biology, yet remains constrained by random insertion, inefficient homologous recombination, and gene targeting. Here, we present PrimeStack, a DSB-independent platform that integrates prime editing with the unidirectional large serine integrase Bxb1, leveraging evolved variants for enhanced activity, to achieve the programmable insertion of multigene, multikilobase cargos at predefined genomic safe harbors in rice. Optimized prime editors first install *attP* landing sites with high fidelity and heritability followed by Bxb1-mediated recombination that generates irreversible integration of genetic information. PrimeStack achieves integration frequencies of approximately 43–46% (as detected by junction-specific PCR in rice calli), with phenotypic neutrality in regenerated plants, comparing favorably with bidirectional Cre-lox systems. We validate its utility by achieving targeted insertion of a carotenoid cassette at an experimentally confirmed genomic safe harbor. PrimeStack delivers a modular, site-specific gene-stacking platform that enables predictable, multigene trait pyramiding and pathway construction at genomic safe harbors, thereby accelerating the development of improved and resilient crop varieties, as well as scalable plant-based biomanufacturing and a powerful chassis for synthetic biology.

**Graphical Abstract:** 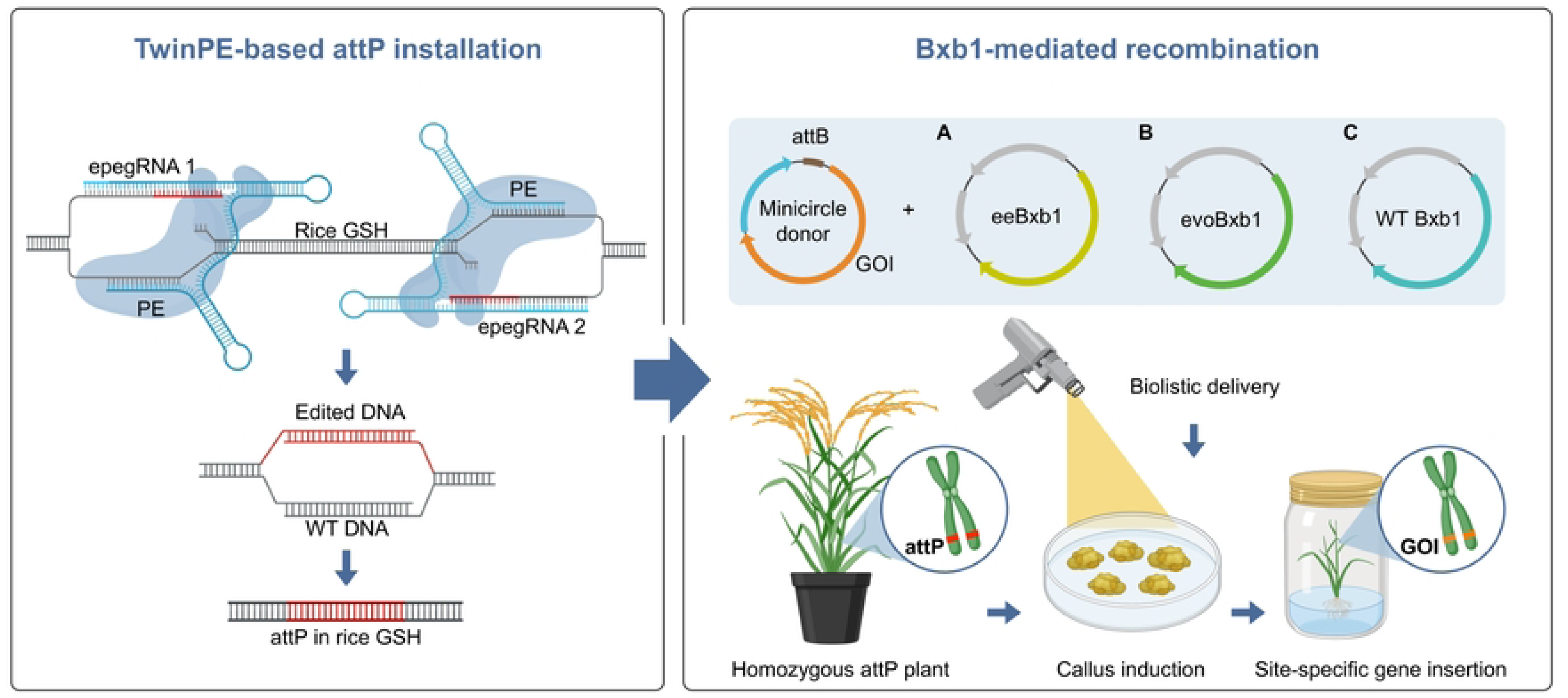

## Introduction

A central challenge in plant biotechnology and synthetic biology is achieving highly efficient, precise, and site-specific integration of multikilobase genetic information into genomic safe harbors (GSH), which could unlock transformative advances in crop improvement and green biomanufacturing [1–4]. Targeted addition of single or multigene traits into predefined, neutral loci promises predictable expression, stable inheritance, and modular engineering, enabling enhanced yield, climate resilience, nutritional biofortification, and the sustainable production of high-value therapeutics, biomaterials, and industrially relevant compounds in plant chassis [5–8]. However, despite decades of progress, achieving high-efficiency, universal, and precise large DNA insertion remains a central bottleneck, limiting the rational redesign of complex traits and the full exploitation of plants as programmable biofactories [9–12].

Transgenesis, pioneered through *Agrobacterium*-mediated T-DNA transfer and biolistic delivery, has revolutionized crop biotechnology by introducing traits of value such as herbicide tolerance, pest resistance, and nutritional enhancements [13–16]. However, the inherent random integration of genetic information frequently results in positional variegation, epigenetic silencing, gene disruption, multiple copy insertions, and reliance on selectable markers, which have contributed to regulatory hurdles and public concerns about transgenic crops, including uncertainty over insertional effects, reliance on antibiotic resistance markers, and the unpredictable nature of random integration events, among multiple other societal and institutional factors [17–22]. Cisgenic and intragenic strategies mitigated some concerns by using compatible genetic elements, but failed to resolve unpredictability and expression instability [23–26].

The advent of sequence-specific nucleases such as meganucleases, zinc-finger nucleases, TALENs, and CRISPR–Cas9 has enabled programmable generation of site-specific double-strand breaks (DSBs), shifting the paradigm toward precision editing [27–33]. CRISPR–Cas systems are highly programmable and efficient for the generation of site-specific DSBs across diverse species, including plants. While DSBs excel at generating knockouts via non-homologous end joining, homology-directed repair for targeted insertions of genetic cargo is notoriously inefficient in plant somatic cells, rendering multikilobase integrations impractical [34–38]. Prime editing (PE), a DSB-independent technology that precisely rewrites genetic information via pegRNAs, has expanded capabilities for base substitutions, indels, and small gene sequence addition in plants [39–43]. However, its cargo size limitation (up to 200 bp) precludes the insertion of full genes or pathways essential for most agronomic and synthetic biology applications [44–46].

Site-specific recombinases (SSRs) offer an alternative for large DNA integration. Tyrosine recombinases (e.g., Cre-lox, FLP-FRT) and serine integrases (e.g., PhiC31, Bxb1) have long been used in plants for marker excision and limited cassette exchange [47–52]. Their utility for *de novo* insertions, however, requires pre-installation of recognition sites, a challenge recently addressed by coupling PE with SSRs. PrimeRoot, for instance, employs PE to insert lox sites followed by Cre-mediated integration of up to 11.1-kb cargos in rice, achieving 6–8% efficiency and marker-free trait delivery [13]. However, Cre’s bidirectional nature, even with pseudo-unidirectional mutants, risks insert instability across generations, while shorter recognition sites raise specificity concerns in repetitive plant genomes [53–55].

Large serine integrases such as Bxb1 provide compelling advantages: strict unidirectionality yielding irreversible *attL*/*attR* hybrids, longer asymmetric sites (38–50 bp) for enhanced fidelity, and established robustness for gene stacking in plants [9, 10, 56–59]. In mammalian cells, prime-editing-assisted site-specific integrase gene editing (PASSIGE) and its evolved eePASSIGE variant harness phage-assisted evolution of Bxb1 to achieve >30% integration of >10-kb sequences, outperforming fusion-based approaches and demonstrating therapeutic potential [12, 14, 60]. Recent megabase-scale chromosome engineering further underscores the power of optimized recombinases combined with scar-free strategies [61].

Here, we introduce PrimeStack, a DSB-free platform that integrates PE with Bxb1, including evolved variants, for programmable, irreversible insertion of multigene cassettes into experimentally validated GSH in rice. By generating recombination-ready germplasm with precisely installed *attP* sites at neutral loci, PrimeStack enables high-efficiency docking of large genetic payloads, resulting in phenotypically neutral lines with stable cassette insertion. This technology overcomes the reversibility and efficiency constraints of Cre-based systems, establishes a universal pipeline for large-scale DNA integration across crops, and paves the way for predictive trait pyramiding, complex pathway assembly, and the realization of plants as scalable, sustainable biofactories [61–64].

## Results

### Selection of genomic safe-harbor loci and prime-editing strategy for Bxb1 attP landing pad installation

Although locus-directed integration has been reported in rice [13], experimentally validated GSH sites remain limited, and the feasibility of installing standardized recombinase landing pads at such loci has not been systematically assessed. We therefore sought to empirically evaluate candidate GSH loci for their ability to tolerate precise installation of a Bxb1 *attP* landing site without detectable fitness penalties, thereby establishing a platform for modular, site-specific transgene integration. We selected eight candidate loci for experimental validation in *Oryza sativa* ssp. *japonica* cv. Nipponbare: six loci nominated from prior computational GSH predictions [13] and two loci within the 25S rDNA region, which has been proposed as a permissive genomic compartment for transgene insertion (Fig 1A) [65]. Non-rDNA candidates were defined using stringent exclusion criteria, prioritizing intergenic regions separated from annotated regulatory and noncoding features (≥5 kb from protein-coding genes and predicted promoters/enhancers, ≥30 kb from microRNA loci, ≥20 kb from lncRNA and tRNA genes, and ≥10 kb from centromeric regions). The final panel comprised two targets within a genomic safe harbor shared across 33 rice genomes on chromosome 1, one target each on chromosomes 3, 6, 7, and 11, and two targets within the 25S rDNA region on chromosome 9 (Fig 1B). To benchmark landing-pad installation in a potentially less “neutral” context, we additionally included two loci proximal to coding genes, near the promoter regions of *OsMOC1* and *OsTT1* [66, 67]. The precise genomic coordinates of each installed *attP* site are provided in S1 Table.

**Fig 1.**
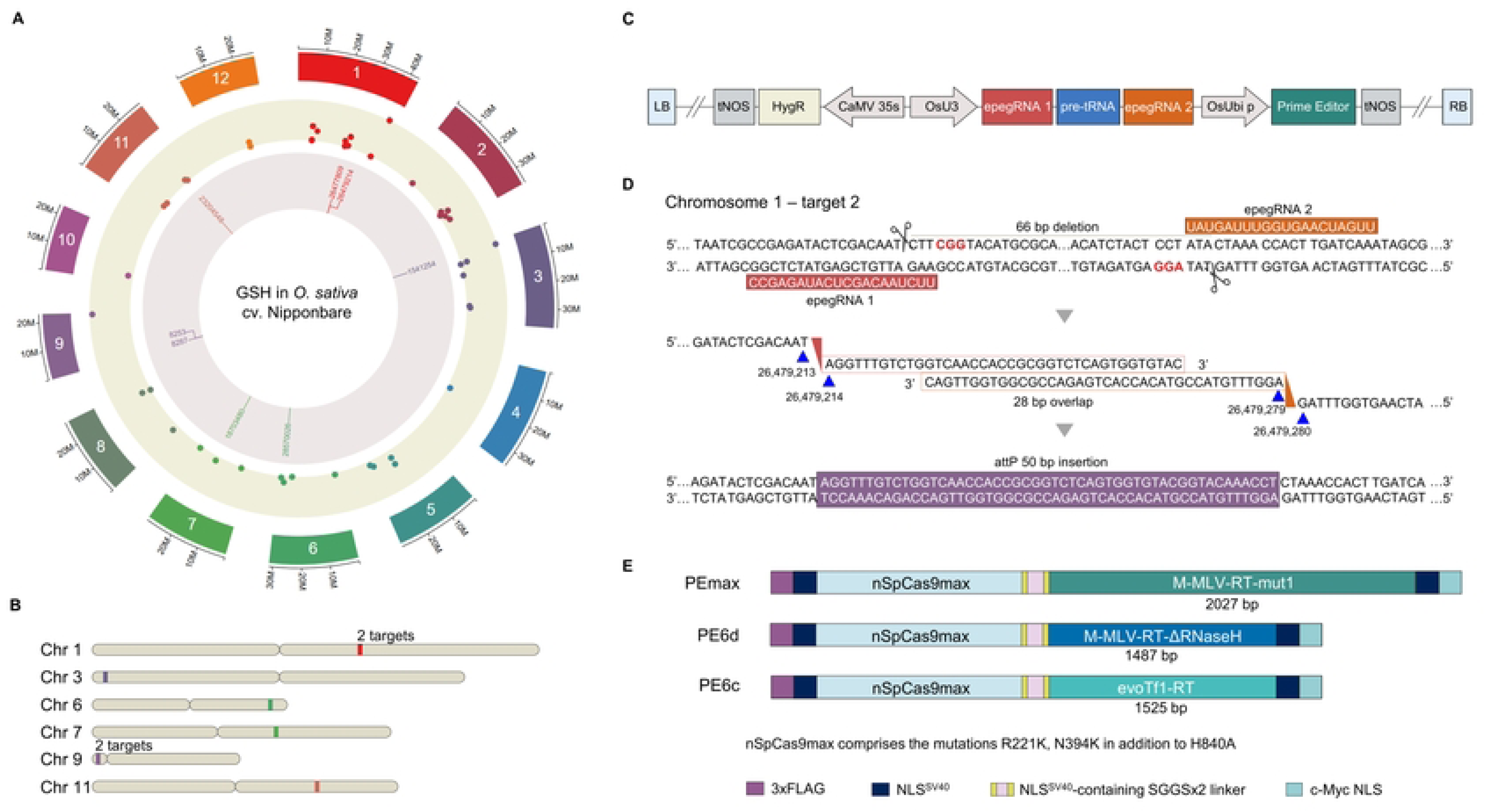
Selection of genomic safe harbors (GSH) and reagents for installation of Bxb1 landing pads (*attP*) in rice genome. (A) Circular ideogram of *O. sativa* ssp. *japonica* cv. Nipponbare genome depicting its relative chromosome sizes (outer ring), the genome-wide distribution of the 44 computationally predicted GSH loci (middle ring), and the genomic positions of the engineered Bxb1 *attP* landing pads introduced at selected GSH (inner ring). Numbers in the inner ring indicate the genomic coordinate of the first nucleotide of each inserted attP site. (B) Chromosomal distribution of GSH loci across individual rice chromosomes. (C) Structure of the *Agrobacterium*-transformation cassette encoding prime-editing components for attP installation. (D) Representative target site Chr 1-2 illustrating Twin-PE-mediated insertion of the Bxb1 attP site. Selected PAM sequences are highlighted in red, and blue arrows indicate the position and orientation of the *attP* insertion relative to the native genome sequence. (E) Overview of the prime editor architectures evaluated in this study.

To install Bxb1 *attP* landing sites, we implemented a dual-pegRNA (Twin-PE) strategy that enables programmable insertion of defined sequences and has been shown to be effective in human cells [7, 12] and plants [13, 46]. For each targeted locus, Twin-PE constructs were delivered into rice callus via *Agrobacterium*-mediated transformation. The T-DNA was designed to express two epegRNAs under the OsU3 promoter and a prime editor under the rice ubiquitin promoter. A hygromycin selection cassette was also used to enable selection of the transformants (Fig 1C). Details about the structure and sequences of the dual-epegRNA cassettes can be found in S4 Fig and S6 and S3 Tables.

Twin-PE utilizes two pegRNAs positioned on opposite DNA strands, each with a reverse-transcription (RT) template that is complementary to the other. The RT templates share a 28-bp overlap, which supports the productive joining of the edited intermediates and the insertion of the full 50-bp *attP* sequence (Fig 1D) [68]. To stabilize pegRNAs and increase editing performance, we appended the tevopreQ1 stabilizing motif to the 3′ end of each pegRNA [68] across the ten target sites, generating engineered pegRNAs (epegRNAs). This architecture enabled the installation of an identical 50-bp *attP* element at each selected locus [7, 12].

Because prime editing performance can be strongly locus- and editor-dependent, we compared three prime editor variants, PEmax, PE6c, and PE6d, which were previously reported to exhibit target-dependent differences in mammalian systems and plants. These editors primarily differ in their reverse transcriptase (RT) modules and overall size. PEmax contains the SpCas9 (R221K, N394K and H840A) nickase fused to the M-MLV-RT-mut1, a human codon-optimized reverse transcriptase of 2,027 bp. In contrast, the PE6d variant incorporates the mutations T128N, V223Y, and D200C within the RT domain and lacks the RNase H module, resulting in a substantially smaller protein with a coding sequence of 1,487 bp. Finally, PE6c carries an evolved Tf1 reverse transcriptase of approximately 1525 bp [69–72]. These architectural differences provided an opportunity to evaluate how RT origin, size, and composition influence Twin-PE performance across target loci (Fig 1E).

### Precise and heritable installation of attP landing pads

Regenerated T0 plants were screened by PCR to identify *attP* insertion events. Among the ten targets evaluated, correct *attP* installation was detected at four genomic safe-harbor loci located in Chr 1-2, Chr 3, Chr 7 and Chr 11, as well as at one site proximal to the promoter region of *OsMOC1*. Fig 2A shows representative events containing the edited amplicons at the engineered GSHs in comparison with the WT counterparts, with expected edited bands indicated by red arrows. Single bands at the expected edited sizes for the Chr 1-2 and Chr11 (263 bp and 194 bp, respectively) evidenced homozygous insertion events. The edited event at Chr 3 displayed two bands, one corresponding to the expected edited amplicon (291 bp) and an additional band of approximately 250 bp that does not correspond to the WT allele, whose expected size is 340 bp. For the selected event at the Chr 7 GSH, three bands were observed, suggesting chimerism.

**Fig 2.**
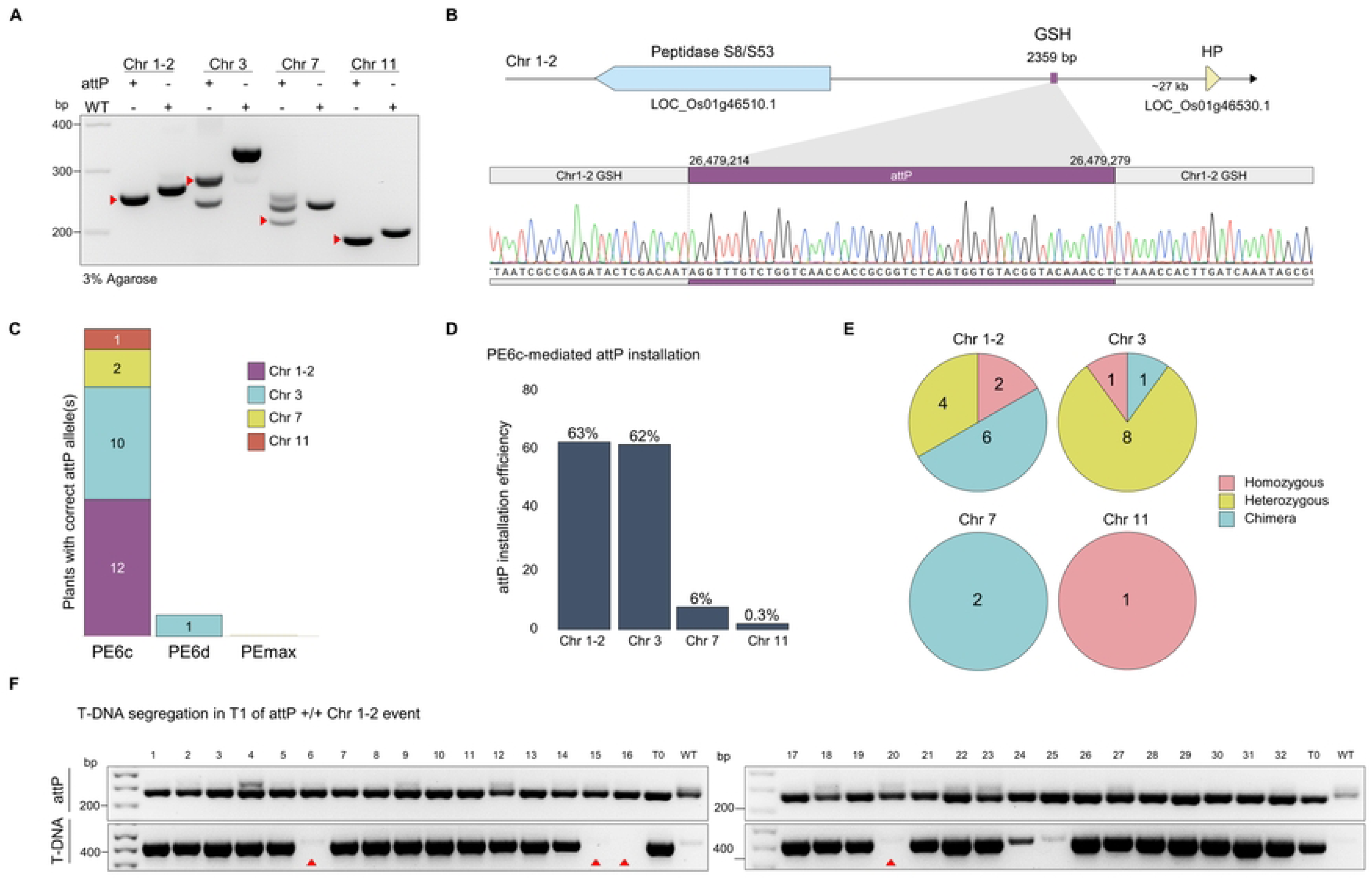
Precise integration and germline transmission of Bxb1 *attP* landing pads. (A) 3% gel electrophoresis confirming successful installation of the *attP* landing pad at selected genomic safe harbor (GSH) loci. Arrowheads indicate the expected amplicons corresponding to correct on-target insertions. (B) Genomic context of the representative *attP* insertion at the chromosome 1-2 GSH, with a Sanger sequencing chromatogram confirming precise on-target integration. (C) Number of regenerated rice plants harboring correctly installed *attP* alleles generated using three prime editor variants (PE6c, PE6d, and PEmax). PE6c produced the highest number of correctly edited plants across multiple loci. (D) Editing efficiency of PE6c-mediated *attP* allele installation across the targeted GSH loci in rice. (E) Zygosity distribution of regenerated plants carrying *attP* insertions at each target site. (F) T-DNA segregation analysis in T1 progeny derived from a homozygous *attP* insertion event at the chromosome 1-2 GSH locus, demonstrating stable *attP* germline transmission and transgene segregation.

To evaluate the fidelity of *attP* landing-pad installation, amplicons from PCR positive T0 plants were subjected to Sanger sequencing. Sequence analysis confirmed the presence of accurate and complete *attP* insertions in the selected events. Fig 2B shows a representative chromatogram demonstrating precise installation of the *attP* site at Chr 1-2.

The majority of the regenerated plants positive for the *attP* insertion were recovered following transformation with the PE6c prime editor. Among loci that yielded detectable insertions, PE6c consistently outperformed PEmax and PE6d, yielding a total of 25 plants carrying the *attP* landing site across four GSH loci. In contrast, PE6d produced only a single regenerated plant with a correctly installed *attP* allele, and no accurate *attP* insertions were recovered using PEmax under our experimental conditions (Fig 2C).

Using PE6c, at least one correctly inserted *attP* allele was detected in 63% of regenerated plants at the GSH in chromosome 1-target 2 and 62% at the chromosome 3 GSH, whereas markedly lower efficiencies were observed at the chromosome 7 GSH (6%) and chromosome 11 GSH (0.3%) (Fig 2D). It should be noted that these percentages represent the proportion of regenerated plants in which any *attP*-containing allele was detectable by PCR and Sanger sequencing, and are therefore qualitative in nature. Because genotyping was performed by conventional PCR rather than amplicon deep sequencing (NGS), the data do not permit per-plant quantification of allele frequency or discrimination between low-level chimerism and stable mono/bi-allelic insertion (Figure 2E). Future studies employing amplicon NGS would provide higher-resolution quantification and enable formal statistical comparison of prime editor variants. At the *OsMOC1* locus, *attP* installation reached 12.5% (S5 Fig). All recovered plants were morphologically normal and produced viable seeds, consistent with phenotypic neutrality of landing-pad installation at these sites under our growth conditions.

Consistent with its higher installation frequency, the chromosome 1-target 2 GSH yielded two independent T0 events that were homozygous for a sequence-accurate *attP* insertion. One of these events was selected for segregation of the T-DNA in the T1 generation. T1 plants were analyzed via PCR and Sanger sequencing to confirm *attP* presence and sequence integrity, while PCR screening for T-DNA enabled identification of T-DNA-free individuals (Fig 2F and S9 Fig). Plants lacking T-DNA were advanced to maturity, and T2 seeds were collected for downstream applications.

### Bxb1-mediated site-specific integration of minicircle donor transgenes in attP-engineered rice calli

Minicircle DNA vectors are minimal, backbone-free circular DNA molecules that are widely used for gene delivery and transgene expression in mammalian systems [12]; however, their application in plant genome engineering remains limited. We therefore explored whether minicircle donors could serve as “clean” payloads for site-specific, recombinase-mediated integration in plants when coupled to Bxb1. To our knowledge, this provides an early demonstration of Bxb1-mediated integration of minicircle donors into engineered *attP* landing pads in the plant genome, offering a potentially streamlined route to precise transgene insertion with minimal vector-derived sequence.

The carotenoid cassette comprises two expression units, each driven by the endosperm-specific glutelin promoter. The first encodes SSU-crtI, consisting of the bacterial *Erwinia uredovora* crtI carotenoid desaturase fused to the chloroplast transit peptide from the pea RUBISCO small subunit (SSU), enabling plastid targeting where carotenoid biosynthesis occurs. The second unit encodes ZmPsy, the maize phytoene synthase that catalyzes the first step of the pathway by converting geranylgeranyl diphosphate (GGPP) into phytoene (Fig 3A).

**Fig 3.**
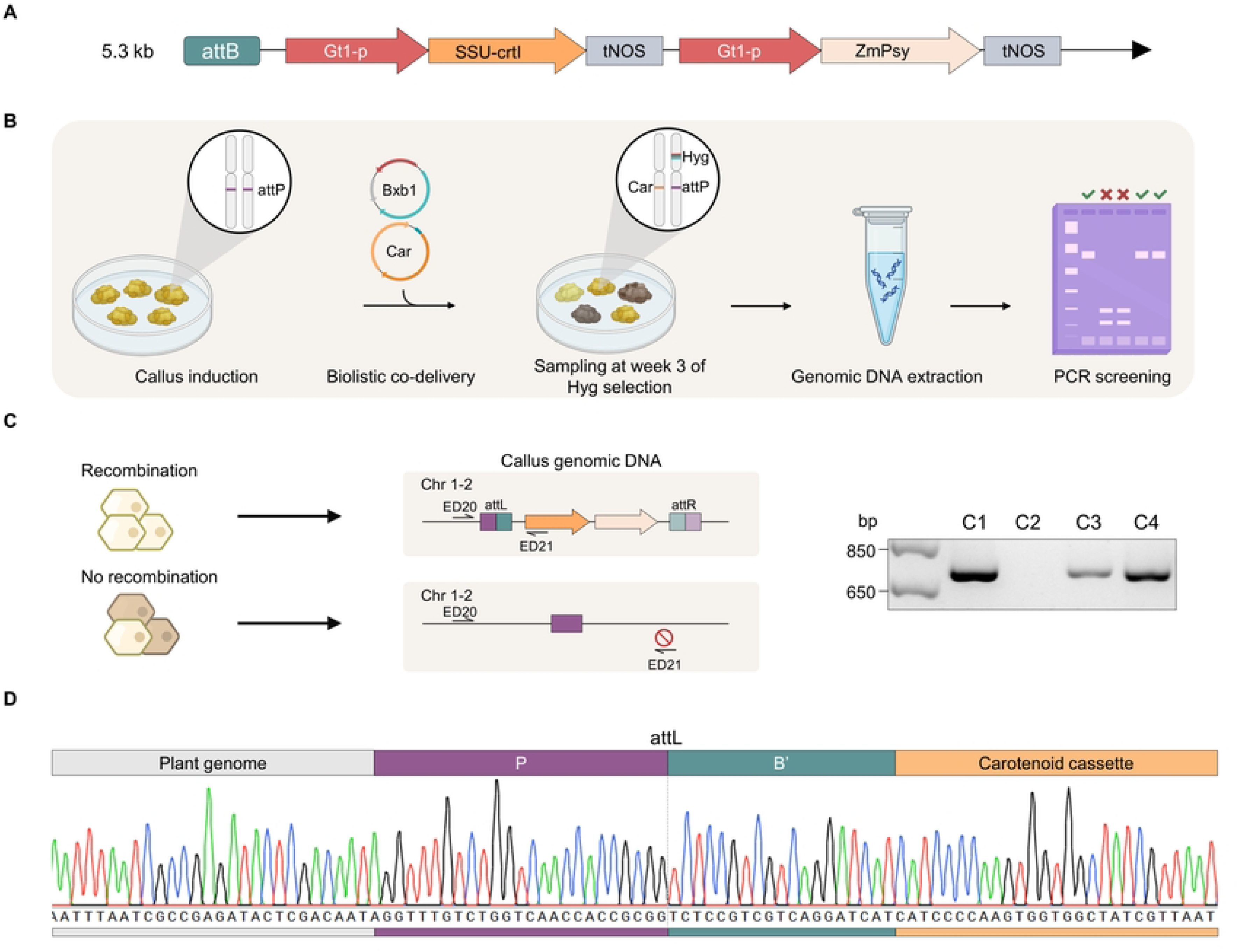
Biolistic co-delivery of a carotenoid minicircle donor and Bxb1 recombinase into *attP*-engineered callus. (A) Schematic of the carotenoid biosynthesis cassette highlighting the attB site and the metabolic pathway components inserted in this study. (B) Transformation workflow: calli were induced from seeds harboring *attP* sites at defined genomic loci. A plant expression vector encoding wild-type (WT) Bxb1 recombinase and a hygromycin resistance marker, together with the minicircle carotenoid donor carrying attB sites, were introduced into engineered calli by particle bombardment. (C) Identification of precise insertion events by junction PCR. (D) Representative Sanger sequencing chromatogram of PCR-positive callus samples confirming correct formation of the attL junction at the targeted integration site.

To test the Bxb1-mediated insertion of the minicircle donor, callus derived from the T-DNA-free homozygous line containing *attP* at the GSH Chr1-2 was subjected to particle bombardment with the purified minicircle donor carrying the Bxb1 *attB* site and the carotenoid biosynthesis cassette (5.3 kb), together with a plasmid expressing the Bxb1 recombinase (Fig 3B). Junction-specific products were detected, demonstrating successful Bxb1-mediated integration of the donor cassette at the intended genomic landing pad (Fig 3C). Sanger sequencing of representative left-junction amplicons verified the presence of the expected *attL* junction and showed that the flanking genomic and donor sequences were accurately preserved without detectable perturbations (Fig 3D).

### Targeted stacking of functional cassettes at a single genomic locus

Having established site-specific integration of a 5.3-kb metabolic cassette, we next challenged our platform with a larger, dual-function payload. We assembled a 9.4-kb cassette combining the carotenoid biosynthesis pathway with a mutated version of the acetolactate synthase gene (*OsALS* W548L), which confers resistance to ALS-inhibiting herbicides such as bispyribac sodium (Fig 4A). T-DNA-free callus derived from the rice line containing *attP* landing pad lines on GSH Chr1-2 was bombarded with the 9.4-kb minicircle donor together with each of the three Bxb1 recombinase variants. After three weeks of hygromycin selection, site-specific integration was assessed by junction PCR. Left junction integration was detected by amplification of an 810-bp product across multiple callus mixtures for all three Bxb1 variants, and right junction amplification yielded products of the expected 502 bp (Fig 4B-D). Sanger sequencing confirmed accurate left and right recombination junctions (Fig 4E-F). Although regeneration of plants carrying the dual-trait donor was not achieved under the tested selection conditions, the robust molecular validation at the callus stage establishes successful site-specific stacking of multiple traits at a single genomic locus.

**Fig 4.**
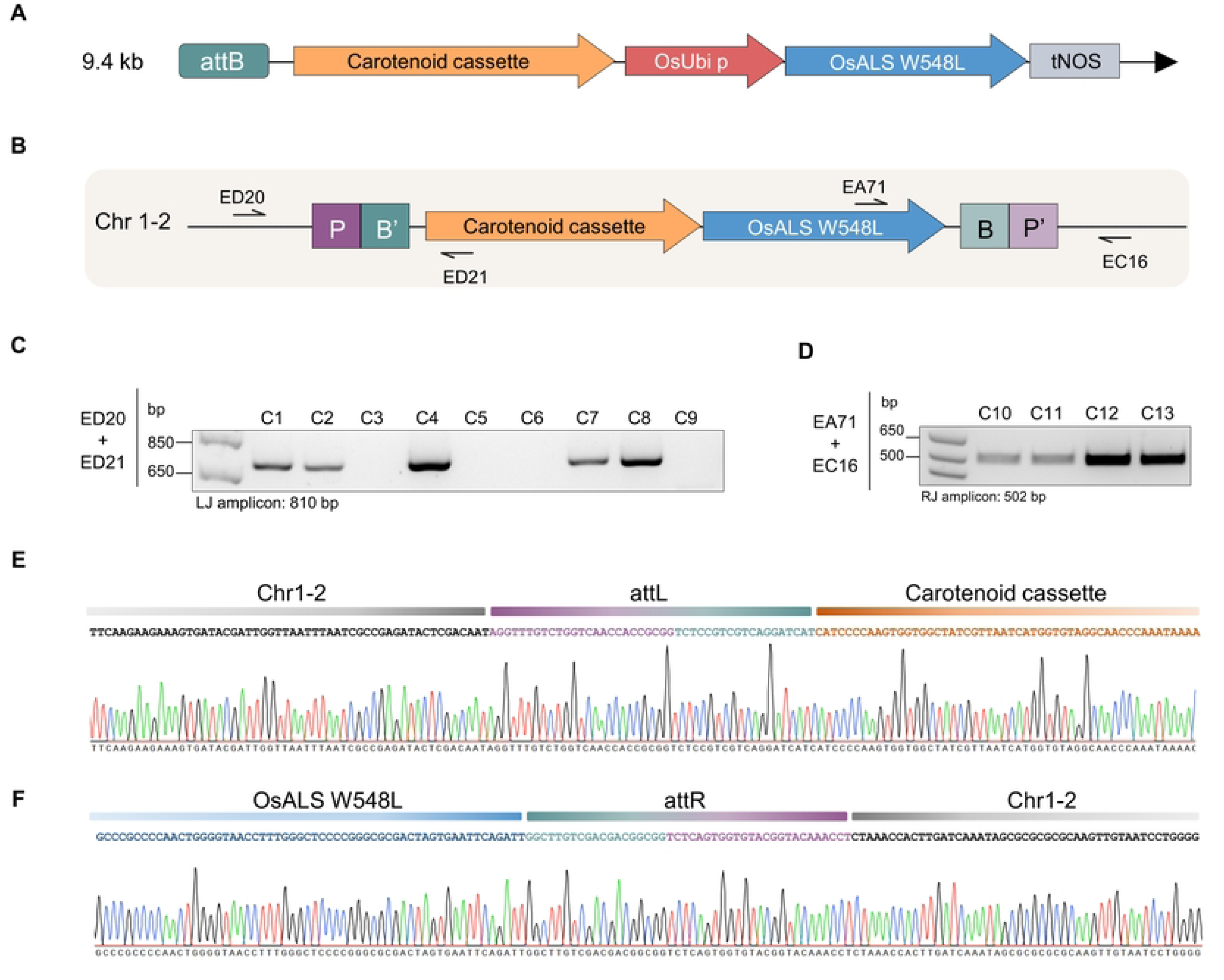
Precise Bxb1-mediated integration of the carotenoid-*OsALS* W548L cassette in callus. (A) Schematic of the carotenoid-*OsALS* W548L cassette used for site-specific integration. (B) Diagram illustrating Bxb1-mediated recombination and targeted insertion of the carotenoid-*OsALS* W548L cassette into the plant genome. (C) Agarose gel electrophoresis of PCR amplicons confirming the presence of the left junction in callus samples. Non-transformed callus (no Bxb1 or donor) was included as a negative control and produced no junction amplicon. (D) Agarose gel electrophoresis confirming the presence of the right junction. Non-transformed callus was included as a negative control. (E) Sanger sequencing chromatograms of the left junction and (F) right junction, confirming precise insertion of the carotenoid-*OsALS* W548L cassette at the Chr 1-2 genomic safe harbor.

### Site-specific integration efficiencies using engineered Bxb1 variants

Recent protein engineering efforts have generated Bxb1 variants with enhanced recombination activity in mammalian cells, including evoBxb1 and eeBxb1 [12]. We hypothesized that these variants could improve integration performance in plant cells. To evaluate this, the carotenoid donor minicircle was co-delivered with WT Bxb1, evoBxb1, or eeBxb1, and integration outcomes were quantified by junction PCR. Both recombination junctions were detected using primer pairs designed to anneal to the carotenoid cassette and the rice Chr1-2 GSH locus (Fig 5A-C). It should be noted that genotyping by junction-specific PCR, including with nested amplification, is inherently biased toward detecting successful integration events and does not capture un-recombined alleles in the bulk genomic DNA. Therefore, the “installation frequency” reported here reflects the proportion of sampled calli/plants in which at least one integration event was detectable by PCR, possibly down to a single template molecule, and is a qualitative rather than quantitative measure of allele-level integration efficiency. Amplicon deep sequencing (NGS) and long-read sequencing (e.g., Oxford Nanopore or PacBio) would provide an unbiased estimate of stable allelic integration frequency and enable formal comparison between Bxb1 variants.

**Fig 5.**
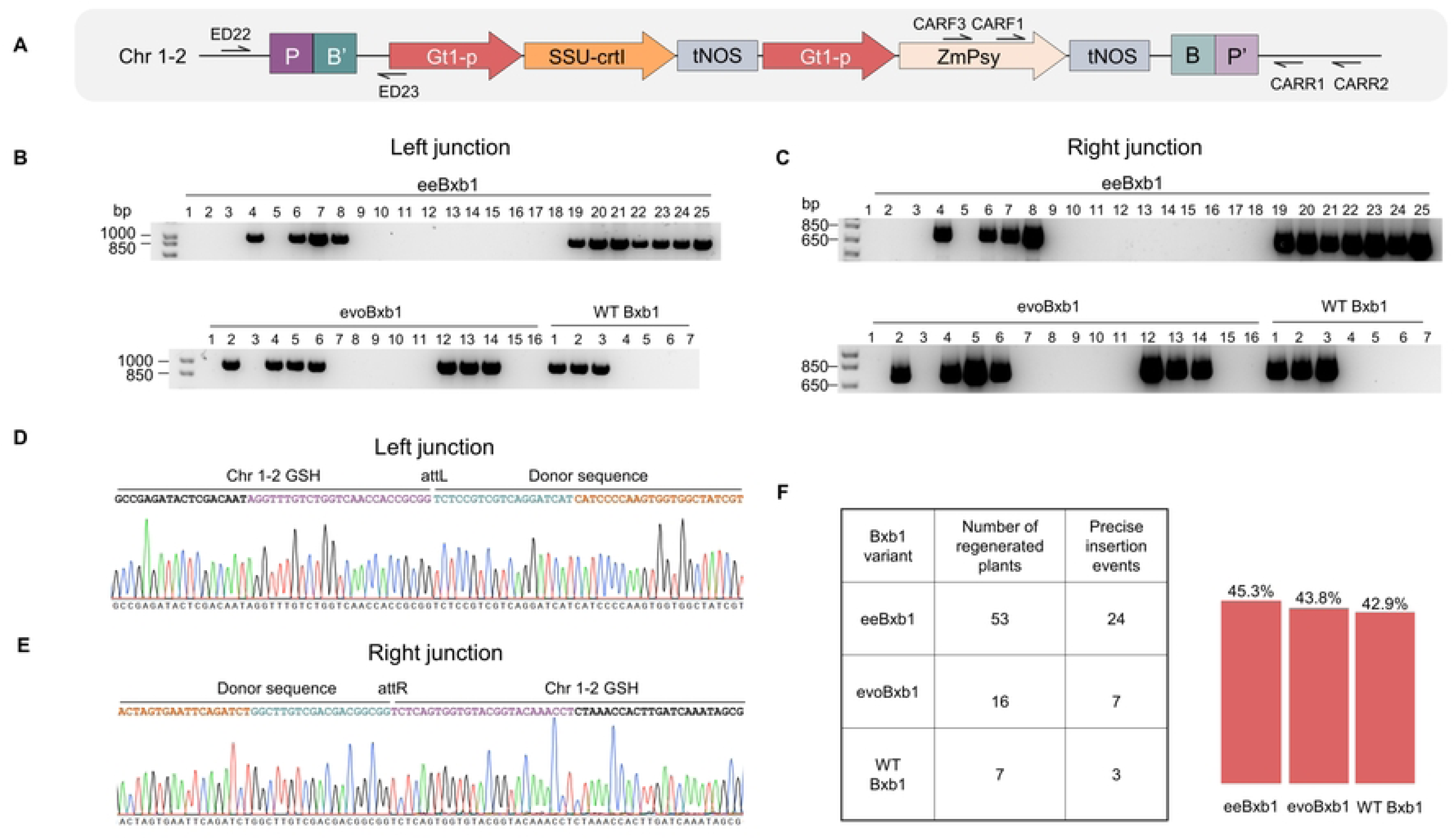
Comparison of Bxb1 recombinase variants for integration of large DNA donors in rice. (A) Schematic of the carotenoid biosynthesis cassette integrated into the plant genome via Bxb1-mediated recombination. Black arrows denote primer pairs used to amplify the left and right junctions. (B) Agarose gel electrophoresis of PCR amplicons confirming the presence of the left junction and (C) the right junction in regenerated plants. Non-transformed callus (no Bxb1 or donor) was included as a negative control in both panels B and C and produced no junction amplicon. In panel C, the right-junction amplicon in some samples was detected only after nested PCR; expected band sizes are indicated by molecular weight markers on the gel. Thick bands in panel C reflect PCR amplification under nested conditions; annotated ladder positions are indicated to assist size estimation. (D) Representative Sanger sequencing chromatograms of the left junction and (E) the right junction, confirming accurate and precise recombination at the attachment sites following transformation with eeBxb1, evoBxb1, or WT Bxb1. (F) Integration efficiency of the carotenoid cassette at the Chr 1-2 GSH, quantified in regenerated plants by junction-specific PCR and validated by Sanger sequencing.

All recovered sequences precisely matched the predicted recombination products generated by Bxb1-mediated integration, confirming accurate site-specific insertion (Fig 5D-E). For the eeBxb1 and minicircle co-delivery, 53 plants carrying validated site-specific cassette insertion were identified. Numerically, eeBxb1 showed the highest integration frequency (45.8%), followed by evoBxb1 (43.7%) and WT Bxb1 (42.8%) (Fig 5F). However, given the sample size (n = 53 for eeBxb1), these differences are modest and should be interpreted with caution, as the confidence intervals for the three variants broadly overlap and no formal statistical test was performed to establish significance. Accordingly, we do not conclude that eeBxb1 is statistically superior to WT Bxb1 under the conditions tested, although the trend toward higher efficiency is consistent with its improved activity in mammalian systems. The limited improvement relative to WT Bxb1 was notably smaller than enhancements reported in human cells [12], suggesting that the advantages conferred by engineered Bxb1 variants may be context dependent and influenced by delivery method, genomic locus, or host-specific constraints.

## Discussion

The precise, site-specific integration of large DNA sequences into plant genomes has long represented a formidable barrier in crop biotechnology and synthetic biology. In this study, we introduce PrimeStack, a pioneering DSB-independent platform that synergistically couples CRISPR-mediated prime editing (PE) with the large serine integrase Bxb1 recombinase to enable programmable, precise insertion of kilobase-scale genetic cargos into GSH. PrimeStack exploits Bxb1’s strictly unidirectional recombination to stack multi-gene cassettes, generating stable *attL*/*attR* hybrids that are inherently resistant to reversal without the need for exogenous factors.

PrimeRoot, the closest prior system, couples enhanced paired prime editors with Cre for DSB-free insertions of up to 11.1 kb in rice, enabling promoter swaps and marker-free disease resistance stacking at efficiencies of 6–8% [13]. However, Cre’s bidirectional nature necessitates transient expression or segregation to avert instability, constraints absent in PrimeStack’s unidirectional framework. Similarly, the mammalian PASSIGE/eePASSIGE system, employing PACE-evolved Bxb1 for 23–30% gene-sized integrations in human cells [12], provides a blueprint for PrimeStack’s enhancements. Evolved and engineered Bxb1 recombinase variants (evoBxb1 and eeBxb1) have previously been reported to mediate up to ∼60% donor integration in human cell lines [12], corresponding to multi-fold improvements over wild-type Bxb1. In contrast, application of the prime editing–Bxb1 system in rice yielded integration frequencies of 42.9–45.3% across Bxb1 variants, with eeBxb1 achieving the highest frequency (45.3%), followed closely by evoBxb1 (43.8%) and wtBxb1 (42.9%) as detected by junction PCR.

Notably, the relative performance differences among Bxb1 variants in plants were modest, in contrast to the more pronounced gains reported in mammalian systems. This suggests that, in plant cells, host-specific constraints, delivery modality (particle bombardment), tissue culture context, and regeneration capacity may play a dominant role in limiting recombination outcomes, thereby reducing the apparent benefit of engineered recombinase variants. Importantly, despite these constraints, our platform supported robust site-specific integration, demonstrating that high-efficiency recombinase-mediated cassette integration is achievable in plants without selection markers.

Several limitations of the current study warrant acknowledgement and point toward key future priorities. First, genotyping was performed exclusively by junction-specific PCR and Sanger sequencing; while this confirms precise recombination at the targeted locus, it cannot exclude additional random integration of minicircle donor DNA elsewhere in the genome via illegitimate recombination. Whole-genome sequencing (WGS) of a representative set of regenerated plants will be required to comprehensively assess integration specificity and exclude off-target insertion events, a particularly important consideration given the use of biolistic delivery, which can introduce multiple DNA copies simultaneously. Second, two of our candidate loci were located within the 25S rDNA region. Because the rice genome harbors hundreds of tandemly repeated rDNA units on chromosome 9, prime editing at these sites may target multiple loci simultaneously, and the precise number and chromosomal identity of targeted copies may vary between experiments. The rDNA-targeted results should therefore be regarded as exploratory; future experiments should incorporate long-read WGS or fiber-FISH to confirm single-locus targeting. Third, from a regulatory standpoint, the Bxb1 *attP* and *attB* recognition sequences used here are 38–50 bp in length. Under proposed EU New Genomic Techniques regulations that classify plants retaining more than 20 consecutive foreign base pairs as conventional GMOs, lines carrying these att sites would fall under full GMO regulatory oversight. Engineering shorter functional att site variants, or adopting recombinase systems with compact recognition sequences compatible with such thresholds, represents an important future direction for translational deployment in regulated markets. Fourth, the current PrimeStack workflow is inherently stepwise: it requires (i) *attP* installation by prime editing with selection and plant regeneration, (ii) T-DNA segregation to recover transgene-free germplasm, and (iii) a second biolistic transformation with the donor and integrase, a process that is labor-intensive and time-consuming. An “all-in-one” co-delivery strategy that simultaneously provides the prime editing machinery, the Bxb1 integrase, and the donor DNA in a single transformation event, as demonstrated for PrimeRoot and the recently described PCE system [14], would substantially streamline the workflow and broaden its practical applicability. Fifth, a direct quantitative comparison of PrimeStack efficiency with Cre-lox-mediated approaches at identical loci and with matched cargo sizes was not performed here; such head-to-head experiments are warranted in follow-up work. Sixth, the recently reported PCE system [14] demonstrates that the bidirectionality limitation of classical Cre-lox can be overcome in plants, further expanding the toolkit for precision plant genome engineering and warranting systematic comparative evaluation against Bxb1-based approaches.

PrimeStack’s emphasis on GSH engineering amplifies its utility. These loci ensure predictable and stable expression without disrupting endogenous functions, mitigating biosafety concerns, and facilitating multiplexed trait stacking for synthetic biology applications, such as biofactories or phytoremediation. PrimeStack offers a paradigm shift toward predictive, design-centric plant engineering. In crop biotechnology, it expedites the prototyping of climate-adaptive varieties by integrating nitrogen-use efficiency modules, carbon-sequestration pathways, or drought-tolerance networks into elite germplasm. Notably, precise GSH-targeted integration of a carotenoid biosynthesis cassette establishes a foundation for carotenoid biofortification in rice, highlighting the potential of PrimeStack for sustainable crop intensification. In plant synthetic biology, PrimeStack provides a powerful chassis for assembling orthogonal circuits, biosensors, and reprogrammed metabolisms at reliable loci. Future iterations, incorporating AI-guided Bxb1 optimizations and expanded GSH repertoires, promise to further amplify its scope, cementing its role in reshaping targeted trait engineering in diverse crop species.

## Materials and methods

### Plant material

Mature dry seeds of *Oryza sativa* ssp. *japonica* cv. Nipponbare were used as explants for callus induction. Seeds were dehusked and surface-sterilized with 70% (v/v) ethanol for 1 minute, followed by incubation in 80% (v/v) commercial bleach solution containing one drop of Tween 20 for 1 hour with continuous shaking. Seeds were rinsed five times with sterile distilled water and placed on 2N6 medium [73] for callus induction.

### General methods

Gibson assembly was used to clone all the prime editors, Bxb1 variants and parental plasmid using NEBuilder HiFi DNA assembly master mix (New England Biolabs). PCR was performed using Phusion High-Fidelity DNA Polymerase (New England Biolabs), Q5 High-Fidelity DNA Polymerase (New England Biolabs) or Phire Plant Direct PCR Master Mix (ThermoFisher). DNA oligonucleotides were obtained from Integrated DNA Technologies (IDT). All PCR products were purified using the QIAquick PCR Purification Kit (QIAGEN). Dual-epegRNAs were obtained from Genscript. All vectors were purified using QIAprep Spin Miniprep kits (Qiagen). All plasmids used in this study are presented in S1 Table. All PCR primers used in this study are presented in S2 Table.

### Prime editor plasmid construction and cloning strategy

DNA fragments encoding prime editors and epegRNAs were cloned into the pCREATE-51 (pC-51) backbone. pC-51 is a modified plasmid derived from pRGEB32 that enables expression of the prime editors PE6c, PE6d, and PEmax under the maize Ubiquitin promoter with a NOS terminator, and the pegRNAs under the OsU3 promoter and the Pol III terminator. The PE6c, PE6d, and PEmax template plasmids were obtained from Addgene (#207853, #207854, and #174820). Ligation of each epegRNA cassette into the corresponding intermediate backbone yielded a total of thirty prime editor constructs, 10 per editor variant, each designed to target the ten selected genomic regions (S1 Fig).

### Bxb1 plasmid cloning

For expression of the Bxb1 variants, the pCREATE-51 plasmid was digested with BsrGI and SacI. Addgene plasmids encoding WT Bxb1, evoBxb1, and eeBxb1 (#222337, #222338, and #222339) were digested with EcoRI, gel-purified, and ligated into the prepared backbone to generate three expression vectors (pC-51-WT-Bxb1, pC-51-evoBxb1 and pC-51-eeBxb1). Each final construct drives expression of the corresponding Bxb1 variant under the rice Ubiquitin promoter with a NOS terminator (S2 Fig).

### Minicircle donor production

The minicircle donors were prepared using the MC-Easy Minicircle DNA Production Kit (System Biosciences, MA925A-1). The parental plasmids were cloned into the *E. coli* ZYCY10P3S2T minicircle producer strain. A single colony was inoculated into 2 mL of LB medium supplemented with 50 μg/mL kanamycin and incubated at 30°C for 4 h. Subsequently, 1 mL of the starter culture was inoculated into 200 mL of TB medium and grown overnight at 30°C (10–12 h) until reaching OD600 4–6. Minicircle induction was initiated by adjusting the culture pH to 7.0 and adding 200 mL of induction medium. Minicircle DNA was purified using the Qiagen Plasmid Maxi kit according to the manufacturer’s instructions. Minicircle quality and integrity were assessed by restriction enzyme digestion and visualization on a 1% agarose gel.

### Agrobacterium-mediated rice transformation for attP installation

A total of thirty plasmids encoding prime editors and pegRNAs were transformed into *A. tumefaciens* strain EHA105 by electroporation. Calli derived from mature seeds were pre-cultured on fresh 2N6 medium at 32°C under continuous light for 3 days and then immersed in the *Agrobacterium* suspension for 2 minutes. Following co-cultivation at 28°C in the dark for 3 days, calli were transferred to selection medium containing 40 μg/mL hygromycin for two weeks, and subsequently moved to second selection medium containing 50 μg/mL hygromycin for 5 days. Resistant calli were transferred to N6RH50 regeneration medium. Regenerated shoots were moved to N6FH50 medium for root development and successfully rooted plantlets were transferred to soil and screened by PCR to identify *attP* insertion events [73].

### Identification of attP insertion events

Genomic DNA was extracted from calli 19 days post-transformation under selection or from regenerated plantlets. Transgenic tissue was PCR-genotyped to identify *attP* insertion events. For edited amplicons differing from wild-type by less than 8 bp, HincII restriction digestion was used to accurately distinguish edited from WT alleles. Amplicons were resolved on 1–3% agarose gel, excised, purified, cloned into the pJET1.2/blunt vector, and submitted for Sanger sequencing. Sequencing chromatograms were aligned to reference sequences using SnapGene Software (GSL Biotech) to confirm precise installation and sequence integrity of the inserted Bxb1 *attP* site. Primer sequences for each target and the expected sizes of WT and edited amplicons are provided in S4 Table.

### Generation of homozygous attP and transgene-free lines

T1 progeny of heterozygous and chimeric T0 events were screened by PCR to identify homozygous *attP* individuals. In parallel, T1 plants were genotyped to identify individuals that had segregated away the T-DNA. Selected T1 plants were grown to maturity, and T2 seeds were collected for subsequent experimental procedures.

### Biolistic co-delivery of minicircle donors and Bxb1 into attP-engineered rice callus

Rice calli harboring Bxb1 *attP* sites were subjected to biolistic co-delivery of minicircle donor DNA and plasmids expressing Bxb1 variants. For each bombardment, 1 μg of minicircle DNA and 1 μg of Bxb1 plasmid were mixed with 50 μL of 0.6 μm gold particles (60 μg/μL), followed by addition of 50 μL of 2.5 M CaCl₂ and 20 μL of 0.1 M spermidine. Bombardment was performed using a Bio-Rad PDS-1000/He particle delivery system equipped with 1100 psi rupture disks. Bombarded calli were transferred to 2NBK medium containing 40 μg/mL hygromycin and incubated at 32°C under continuous light for two weeks, then moved to nN6C selection medium with 50 μg/mL hygromycin for 5 days before transfer to R8 regeneration medium [73–75]. Regenerated plants were screened by PCR for site-specific integration.

### Analysis of Bxb1-mediated long DNA fragment integration

Genomic DNA was extracted from calli 19 days post-transformation under selection or from regenerated plantlets. The left and right genome-donor junctions were PCR amplified and resolved by 1% agarose gel electrophoresis, excised, purified, cloned into the pJET1.2/blunt vector, and submitted for Sanger sequencing. Sequencing chromatograms were aligned to reference sequences using SnapGene Software (GSL Biotech; www.snapgene.com) to confirm junction accuracy and detect potential sequence alterations.

### Genomic DNA extraction from plant tissues

Frozen plant leaves were ground to a fine powder, whereas callus tissue was homogenized to release cellular contents. Samples were resuspended in extraction buffer (0.1 M Tris-HCl pH 8.0, 1 mM EDTA, 0.1 M NaCl, 0.1 M LiCl, 0.1 M β-mercaptoethanol, 0.4% [w/v] RNase I) at a ratio of 500 μL per 300 μL of leaf tissue or five callus pieces. Samples were subjected to phenol:chloroform:isoamyl alcohol extraction, and DNA was precipitated with absolute ethanol and 30 μL of 3 M sodium acetate (pH 5.2). Pellets were washed twice with 70% ethanol, air-dried, and resuspended in nuclease-free water. DNA concentration was assessed by NanoDrop UV/Vis spectrophotometry.

## Acknowledgments

We thank members of the Laboratory for Genome Engineering and Synthetic Biology for discussions and help. We thank Professor Pamela Ronald for generously sharing the pAcc-B plasmid used in this study.

## Author contributions

E.S., K.S., H.B. and M.M. designed the research; E.S. and K.S. performed research; E.S., K.S., H.B. and M.M. analyzed the data; E.S., K.S. and M.M. wrote the paper.

## Competing interests

The authors have declared that no competing interests exist.

## Funding

This work was supported by BAS/1/1035-01-01 baseline funding to M.M. The funders had no role in study design, data collection and analysis, decision to publish, or preparation of the manuscript.

## Data availability

All data underlying this article are available in the article and in its online Supporting Information files.

## Supporting information

**S1 Fig.** Prime editor constructs for *attP* installation at ten genomic targets. Schematic representation of the pCREATE-51-based constructs encoding PE6c, PE6d, and PEmax prime editors with dual-epegRNA cassettes targeting each of the ten selected loci. Expression cassettes for the prime editor protein and dual epegRNAs are shown.

**S2 Fig.** Bxb1 expression vector maps. Schematics of the pCREATE-51-based constructs encoding WT Bxb1, evoBxb1, and eeBxb1 recombinases under the rice Ubiquitin promoter.

**S3 Fig.** Cloning strategy overview for parental plasmids. Schematic depicting the sequential assembly steps for PP-Carotenoid, PP-*OsALS* W548L, and PP-Carotenoid-*OsALS* W548L parental constructs.

**S4 Fig.** Dual-epegRNA cassette architecture. Structure and sequence details for the dual-epegRNA cassettes used for *attP* installation, including spacer sequences, PBS lengths, RT template sequences, and tevopreQ1 stabilizing motif positions for each of the ten target loci.

**S5 Fig.** *attP* installation at the *OsMOC1* promoter locus. (A) PCR results showing *attP*-containing amplicons at the *OsMOC1* locus. (B) Sanger sequencing chromatogram confirming precise *attP* insertion near the *OsMOC1* promoter. (C) Installation efficiency at the *OsMOC1* locus. (D) T-DNA segregation analysis in T1 progeny derived from the *attP*-*OsMOC1* event.

**S6 Fig.** Sanger sequencing chromatograms confirming *attP* insertion at Chr 3, Chr 7, and Chr 11 GSHs. Representative chromatograms showing accurate and complete *attP* insertion sequences at each locus.

**S7 Fig.** Characterization of aberrant amplicons at Chr 3 and Chr 7. Sanger sequencing chromatograms of amplicons deviating from expected size at the Chr 3 and Chr 7 GSHs, indicating defective or rearranged versions of edited or wild-type alleles.

**S8 Fig.** Sequence analysis of *OsTT1* candidate events. Sanger sequencing data from six candidate homozygous events at the *OsTT1* locus showing a 15-bp deletion within the *attP* sequence.

**S9 Fig.** T-DNA segregation analysis at Chr 1-2. PCR-based genotyping of T1 progeny from the homozygous *attP* Chr 1-2 event showing T-DNA segregation and identification of T-DNA-free individuals.

**S10 Fig.** T1 segregation analysis at additional loci. (A) PCR screening of T1 progeny from *attP*-Chr 11 event. (B-C) Genotyping of heterozygous and chimeric T0 events advanced to T1. (D) Confirmation of T-DNA retention in T1 plants.

**S11 Fig.** Sequence integrity of *attP* in T1 homozygotes. Sanger sequencing confirming accurate *attP* sequence in T1 homozygous lines derived from heterozygous T0 events.

**S12 Fig.** Minicircle production and validation. (A) Parental plasmid structure for carotenoid minicircle production. (B) Minicircle induction protocol overview. (C-D) Restriction enzyme digestion confirming identity of parental plasmid and resulting minicircle. (E-F) Bxb1 expression vector schematics.

**S13 Fig.** Characterization of the carotenoid-*OsALS* W548L parental plasmid and minicircle donor. Restriction enzyme digestion profiles confirming the identity of the 13.6-kb parental plasmid and the 9.4-kb minicircle donor.

**S14 Fig.** Junction PCR results for the full set of eeBxb1-transformed samples. Gel images and tabulated results for all 53 validated site-specific insertion events identified by left- and right-junction PCR following co-delivery of eeBxb1 and the carotenoid minicircle donor.

**S1 Table.** List of all plasmids used in this study.

**S2 Table.** List of all PCR primers used in this study.

**S3 Table.** List of synthesized dual-epegRNA sequences used for *attP* installation.

**S4 Table.** Primer sequences for genotyping each target locus and expected WT and edited amplicon sizes.

**S5 Table.** Precise genomic coordinates of each installed *attP* site.

**S6 Table.** Sequences of dual-epegRNA cassettes for all ten targets.

## Notes

### Competing Interest Statement

The authors have declared no competing interest.

